# Improved *KCNQ2* gene missense variant interpretation with artificial intelligence

**DOI:** 10.1101/2022.10.20.513007

**Authors:** Alba Saez-Matia, Arantza Muguruza-Montero, Sara M-Alicante, Eider Núñez, Rafael Ramis, Óscar R. Ballesteros, Markel G Ibarluzea, Carmen Fons, Aritz Leonardo, Aitor Bergara, Alvaro Villarroel

## Abstract

Advances in DNA sequencing technologies have revolutionized rare disease diagnosis, resulting in an increasing volume of available genomic data. Despite this wealth of information and improved procedures to combine data from various sources, identifying the pathogenic causal variants and distinguishing between severe and benign variants remains a key challenge. Mutations in the K_v_7.2 voltage-gated potassium channel gene (*KCNQ2*) have been linked to different subtypes of epilepsies, such as benign familial neonatal epilepsy (BFNE) and epileptic encephalopathy (EE). To date, there is a wide variety of genome-wide computational tools aiming at predicting the pathogenicity of variants. However, previous reports suggest that these genome-wide tools have limited applicability to the *KCNQ2* gene related diseases due to overestimation of deleterious mutations and failure to correctly identify benign variants, being, therefore, of limited use in clinical practice. In this work, we found that combining readily available features, such as AlphaFold structural information, Missense Tolerance Ratio (MTR) and other commonly used protein descriptors, provides foundations to build reliable gene-specific machine learning ensemble models. Here, we present a transferable methodology able to accurately predict the pathogenicity of *KCNQ2* missense variants with unprecedented sensitivity and specificity scores above 90%.

## INTRODUCTION

Over the past decade, the rapid progress in genomic technologies has made research and clinical sequencing increasingly accessible [Traynelis *et al*., 2017]. However, identifying the true phenotypically causal variants remains a major challenge, mainly in terms of time and costs [McInnes *et al*., 2021; Finn *et al*., 2009]. In addition, weak effect sizes and correlations among neighboring variants, locus heterogeneity and limitations in sample or pedigree sizes limit the discovery power and resolution of purely genetic studies [Cooper & Shendure, 2011]. This problem is exacerbated for rare diseases and complex pathologies where genetic information by itself is often insufficient [McInnes *et al*., 2021; Mayer *et al*., 2011; MacArthur & Smith, 2010]. This scenario is especially visible for *KCNQ2* gene related pathologies.

*KCNQ2* gene encodes the voltage-gated potassium channel subunit K_v_7.2, which plays a crucial role in neuronal excitability by mediating the M-current [Etxeberrria *et al*., 2008; Schwake *et al*., 2006; Gutman *et al*., 2003; Marrion, 1997]. Initially, autosomal dominantly inherited *KCNQ2* mutations were linked to benign familial neonatal epilepsy (BFNE) [Singh *et al*., 1998; Biervert *et al*., 1998]. Later on, *de novo KCNQ2* mutations were identified in patients with neonatal developmental and epileptic encephalopathy (DEE, EE) [Goto *et al*., 2021; Millichap *et al*., 2016; Pisano *et al*., 2015; Numis *et al*., 2014; Kato *et al*., 2013; Weckhuysen *et al*., 2013; Weckhuysen *et al*., 2012]. *KCNQ2* encephalopathy presents early neonatal onset intractable seizures, usually tonic seizures with suppression burst EEG pattern and severe developmental delay [Millichap *et al*., 2016; Pisano *et al*., 2015; Kato *et al*., 2013; Weckhuysen *et al*., 2013]. Intractable seizures and poor prognosis warrant an urgent need for effective early diagnosis and treatment [Kim *et al*., 2021]. We strongly believe that this effective treatment must start with a correct identification of *KCNQ2* pathogenic variants.

To achieve that aim, the use of computational tools has been extended to many fields of medical sciences, including development of predictive software to assess the pathogenicity of protein variants [Wainberg *et al*., 2018; Thusberg *et al*., 2011]. To date, there are a wide variety of computational tools aiming at predicting the pathogenicity of variants [Katsonis *et al*., 2021]. Some commonly used tools include Sorting Intolerant From Tolerant (SIFT, [Ng, 2003]), Protein Variation Effect Analyzer (PROVEAN, [Choi *et al*., 2012]) and Functional Analysis through Hidden Markov Models (FATHMM, [Shihab *et al*., 2013]), which use a sequence homology-based method; Polymorphism Phenotyping v2 (PolyPhen2, [Adzhubei *et al*., 2010]), which utilizes a sequence and structure-based approach; MutPred2 [Pejaver *et al*., 2020], which combines evolutionary, structural and function-based features; and consensus deleteriousness score (CONDEL, [González-Pérez & López-Bigas, 2011]) which uses a consensus-based approach. Despite relative success in differentiating between severe disease-causing and benign variants in some genes, these tools are not consistently effective in their predictions and their accuracy has been shown to be gene-dependent [Leong *et al*., 2015; Chun & Fay, 2009]. Consequently, there is no *in silico* tool capable of effectively predicting the pathogenicity of *KCNQ2* variants to date with sensitivity/specificity scores above 90%, which characterize a clinical relevant tool [Richards *et al*., 2015]. Nevertheless, machine learning-based techniques have already proven their utility for other diseases related to genes encoding ion channels [Xenakis *et al*., 2021, Sallah *et al*., 2020, Li *et al*., 2017].

Machine learning (ML) refers broadly to the process of fitting predictive models to data or of identifying informative groupings within data. The field of ML essentially attempts to approximate or imitate the ability to recognize patterns of humans, albeit in an objective manner, through algorithms and computation [Greener *et al*., 2021]. Supervised ML refers to techniques in which a model is trained on a range of inputs (or features) which are associated with a known outcome. In medicine, this might represent training a model to relate the characteristics of an individual (e.g., height, weight, smoking status) to a certain outcome (onset of diabetes within five years, for example). Once the algorithm is successfully trained, it should be capable of making outcome predictions when applied to new data [Jenni *et al*., 2019].

It is accepted that the function of proteins emanates from the structure, which in turn depends on its amino acid sequence. However, the use of 3D information has been hampered by the lack of resolved structures at atomic resolution, especially for membrane proteins such as ion channels. The cryo-EM revolution has ameliorated this limitation to great extent, but most of the published structures only provide partial coverage (i.e., some large stretches of amino acids are missing in the coordinate files). For instance, the recent K_V_7.2 structure submitted in the RCSB Protein Data Bank (https://www.rcsb.org/) (PDB: 7CR3) covers only 40% of the total amino acid sequence. Fortunately, the release of AlphaFold2 [Jumper *et al*., 2021] opened up a new era in the understanding of protein structure. The deep neural network of this algorithm, which combines features derived from homologous templates and from multiple sequence alignment to generate the predicted structure, has shown an outstanding accuracy in predicting the three-dimensional structure of proteins with otherwise unknown fold [David *et al*., 2022]. In this study, we investigated methods to capture this structural information to help discriminating the pathogenic consequences of missense mutations. Combining these with more traditional descriptors, we developed the first *KCNQ2-specific* machine learning-ensemble algorithm (MLe-KCNQ2) capable of accurately predicting the pathogenicity of channel K_v_7.2 variants. This novel algorithm, with a sensitivity of 91.75% and a specificity of 90.21%, shows a high accuracy at predicting both pathogenic and benign variants, respectively. Therefore, MLe-KCNQ2 could help clinicians and researchers to interpret the effect of missense mutations in the *KCNQ2* gene as well as facilitate the clinical diagnosis and early choice of personalized therapies for developmental *KCNQ2*-related disorders. The methodology employed here can be extended to the prediction of the pathogenicity of variants occurring in other genes.

## METHODS

### DATASET

Isoform 4 of the KCNQ2 gene encodes a transcript with 872 codons, which could lead to 7.582 single base substitutions, leading to 5.972 single amino acid missense variants (excluding stop codons). To date, more than 1.200 *KCNQ2* variants have been annotated in the ClinVar database (https://www.ncbi.nlm.nih.gov/clinvar/) (16/09/21). These variants are divided into 6 subclasses according to the type of the mutation: frameshift, missense, nonsense, splice site, non-coding RNA, and untranslated region (UTR). In order to develop an efficient classification algorithm, three aspects must be taken into account: (i) each subclass must be analyzed individually because a given feature could be important for one subclass but not relevant for others [Larrea *et al*., 2021]; the chosen subclass must have (ii) an adequate number of variants and (iii) at least two clinical conditions with distinct effect (e.g., benign and pathogenic significance). This study focuses on missense variants because they are the only ones that meet all these criteria.

More than 500 missense variants are already annotated in ClinVar, which are distributed according to their pathogenicity as follows: 6 benign, 12 likely benign, 254 of uncertain significance, 39 of conflictive interpretation, 122 likely pathogenic and 141 pathogenic. It should be noted that some of these variants have been included into more than one subclass. In order to simplify the classification as a binary problem, benign and likely benign variants were grouped into the benign class, and pathogenic and likely pathogenic variants into the pathogenic class. At this point, uncertain significance and conflicting interpretation variants were isolated. As a result, an initial dataset consisting of 12 benign and 240 pathogenic variants was obtained.

In addition, an exhaustive database and bibliographic search was carried out. This procedure allowed us to incorporate missense variants not present in the initial dataset. If a variant was shared between two databases, the most recent entry was chosen. As a result, the number of variants with a reliable diagnosis was increased. In sum, the final dataset consisted of 39 benign and 314 pathogenic variants distributed along all K_v_7.2 functional domains (Figure 1). After that, clinical labels were binarised, with benign and pathogenic variants equal to 0 and 1, respectively.

**Figure 1.**
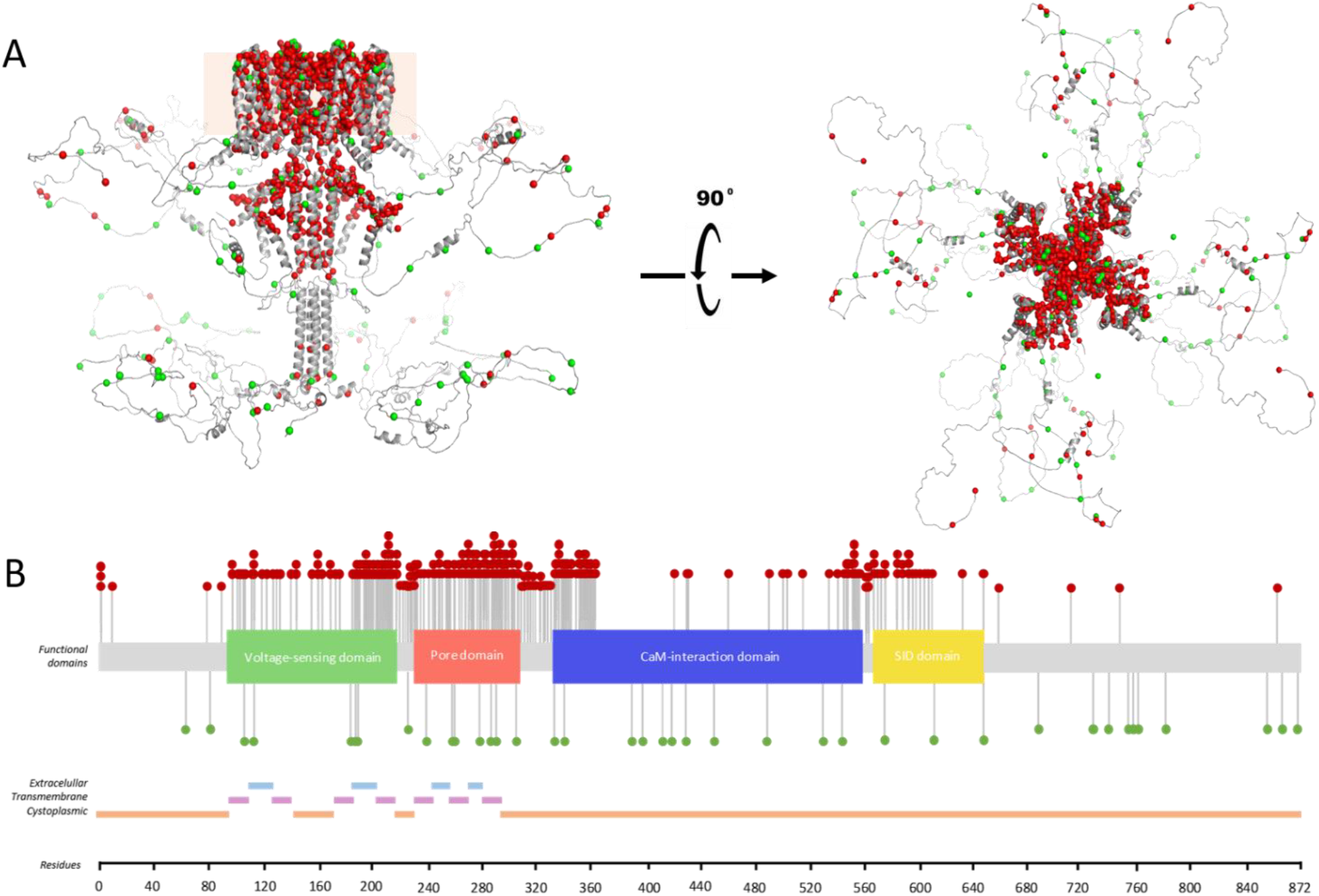
Graphical distribution of the 353 collected variants. **(A)** Distribution of the collected variants on the three-dimensional structure of a K_v_7.2 channel tetramer, lateral (left) and upper view (right). The structure of the K_v_7.2 monomer was built using the best prediction obtained with AlphaFold2. Intrinsically Disordered Regions (IDRs) were modeled using SwissModel [Waterhouse *et al*., 2018] to avoid wrong location of these regions in the membrane and clashes after tetramerization. Both figures were created using PyMol [DeLano, 2002] and VMD [Humphrey *et al*., 1996]. Pathogenic and benign variants are colored in red and green, respectively, and the membrane is indicated with an orange square in the lateral view. **(B)** Distribution of the collected variants along the main functional domains of the channel. Both the 305 pathogenic variants and the 39 benign variants are distributed throughout the entire K_V_7.2 sequence using the previous color code. In addition to being predominant, pathogenic variants are found most abundantly in the voltage-sensing domain (VSD), the pore and the first residues of the CaM-interaction domain. In contrast, the collected benign variants are homogeneously distributed. Information about extracellular (light blue), transmembrane (pink) and cytoplasmic (orange) segments is also shown.

### VARIANTS CHARACTERIZATION

The final dataset was characterized using 16 features. These characteristics can be divided into 3 different subgroups: features related to physico-chemical changes (amino acid charge, hydrophobicity, molecular weight, volume, polarity, aromaticity and amino acid mean solvent accessibility), biological- evolutive features (evolutive conservation value across species of the substituted residue, original and mutated residue, topological and functional affected domains, and Missense Tolerance Ratio or MTR score [Traynelis *et al*., 2017]); and structural features (secondary structure affected by the mutation, AlphaFold2 pLDDT score and location within the structural landscape of *KCNQ2*).

### MACHINE LEARNING ALGORITHMS

The different procedures shown below were performed using Python (v3.8.8) [Van Rossum & Drake, 2009] and the pandas (v1.2.4) [McKinney, 2010], NumPy (v1.20.1) [Harris *et al*., 2020] and scikit-learn (v1.1.1) [Pedregosa *et al*., 2011] libraries. Visualizations were performed using the Matplotlib library (v3.3.4) [Hunter, 2007].

### DATA PREPROCESSING

The final dataset was subjected to format homogenization, control of outliers and missing values as well as elimination of duplicates. As a result, 9 pathogenic variants were finally discarded. The remaining 344 variants were randomly assigned to training (75%) and test (25%) sets. The training and test sets consisted of 258 (31 benign and 227 pathogenic) and 86 variants (8 benign and 78 pathogenic), respectively. As the dataset was clearly unbalanced, an oversampling technique was applied in the training set in order to artificially increase the number of benign variants. Oversampling was performed by increasing the amount of minority class samples by producing new instances or repeating some of them [Mohammed *et al*., 2020]. In sum, 113 benign variants were included. After that, categorical features (e.g. amino acid charge change or topological affected domains) were encoded. First, these categorical values were transformed into numerical labels using the scikit-learn LabelEncoder class. After that, a one-hot encoding scheme was applied (scikit-learn OneHotEncoder class).

### MODEL DEFINITION, TRAINING AND OPTIMIZATION

In a first approach, three simple ML algorithms were trained: Ridge Regression (i.e., a linear regression model that implements L2 norm for regularization) (LR2), Support Vector Classifier (SVC) and Random Forest (RF). Previous studies have shown that these models are useful in variant pathogenicity prediction and also in unbalanced class scenarios [Greener *et al*., 2021]. Each algorithm was trained individually using a pipeline with data standardization and the classifier. A feature selection method based on F-score (ANOVA) was applied during training. This feature selection method is used to select a subset of relevant features for effective classification, i.e., to automatically identify the most informative features out of the original ones and use them in the classification problem [Hoque *et al*., 2014, Polat *et al*., 2009]. Hyperparameters of each model were adjusted using a Grid Search algorithm on a 10 times 10-fold cross validation. A random seed was used to ensure reproducibility.

In order to improve the overall efficiency and reduce bias of individual models, an ensemble algorithm was developed (MLe-KCNQ2). This MLe-KCNQ2 algorithm consists of the previous LR2, SVC and RF models and predicts on the basis of aggregating the findings of each individual estimator in a soft voting configuration, i.e., on the argmax of the sums of the predicted probabilities [Pedregosa *et al*., 2011] (Figure 2).

**Figure 2.**
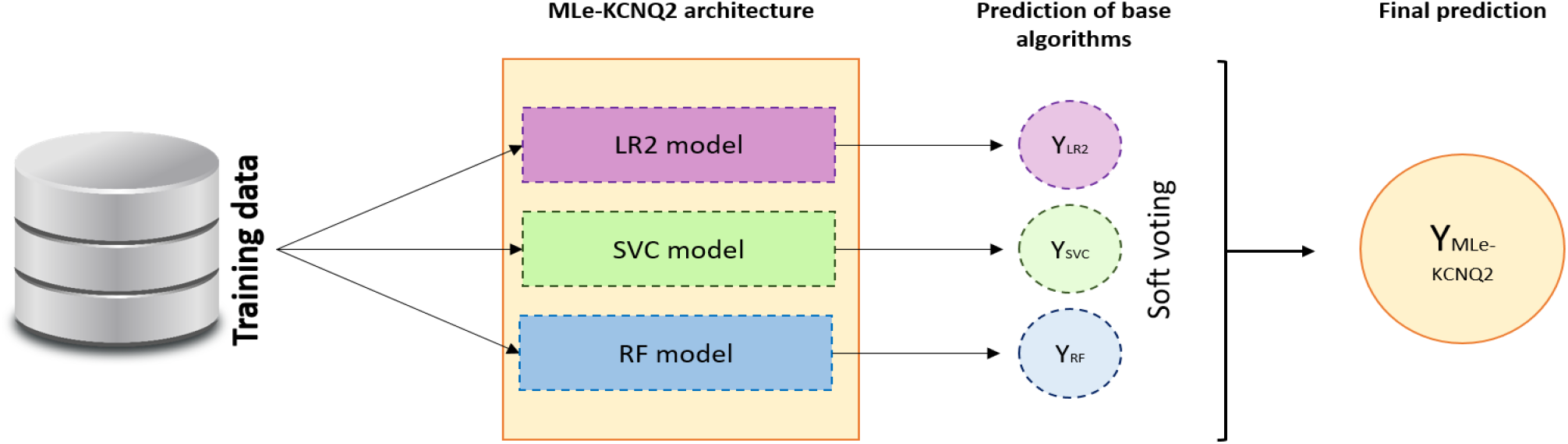
Architecture of MLe-KCNQ2 algorithm. MLe-KCNQ2 is an ensemble algorithm consisting of three individual estimators: a Ridge Regression (LR2), a Support Vector Classifier (SVC) and a Random Forest (RF). The final prediction of the MLe- KCNQ2 algorithm was obtained by combining the individual predictions of the base estimators in a soft voting configuration.

### MODEL EVALUATION

Models were evaluated by estimating sensitivity and specificity values from confusion matrices. Areas under the Receiver Operating Characteristic (ROC) curves or AUC-ROC were also analyzed. All of these metrics are useful in class imbalance scenarios and are clinically relevant [Japkowicz, 2013]. On the one hand, sensitivity is the metric that evaluates the algorithm’s ability to correctly predict true positives (see Equation 1). In the particular case of this study, it would be the ability of the model to distinguish pathogenic variants. On the other hand, specificity is the metric that assesses the ability of the algorithm to correctly differentiate true negatives (see Equation 2). In this context, the ability of the model to distinguish benign variants.

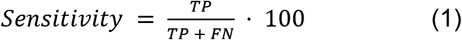

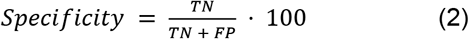

where TP, FP, TN, FN are the true positives, false positives, true negatives and false negatives, respectively.

The ROC curve expresses the relationship between sensitivity and (1 – specificity) for different classification thresholds. The AUC-ROC would be the numerical value that summarizes the predictive ability of that model and is obtained by integrating the area under the ROC curve [Muschelli, 2019]. The AUC-ROC can take any value between 0 and 1 and it is a good indicator of the goodness of the model [Kim *et al*., 2017]. In general, an AUC-ROC of 0.5 suggests that the model has no discrimination (i.e., would not distinguish between benign and pathogenic variants), 0.7 to 0.8 is considered acceptable, 0.8 to 0.9 is considered excellent, and above 0.9 is considered outstanding [Hosmer, 2013].

### COMPARISON TO ALTERNATIVE APPROACHES

The best model was then compared to some commonly used online predictive tools such as FATHMM (http://fathmm.biocompute.org.uk/disease.html), CONDEL (http://bbglab.irbbarcelona.org/fannsdb/query/condel), PolyPhen2 (http://genetics.bwh.harvard.edu/pph2/bgi.shtml), MutPred2 (http://mutpred.mutdb.org/#qform), SIFT (https://sift.bii.a-star.edu.sg/) and PROVEAN (http://provean.jcvi.org/index.php). All collected variants were used in this comparative analysis.

### MODEL PREDICTION

Finally, the model was used to predict the pathogenicity of 293 uncertain significance or conflicting interpretation variants annotated in ClinVar that were not part of the initial dataset. As 3 of these variants were included in both subclasses, only one of them was taken into account. As a result, 290 variants were predicted. In addition to predicting the clinical label, the confidence of each prediction was estimated. A subset of predicted variants was then selected for a biological interpretation, based on the prediction confidence and the availability of electrophysiological studies already reported in the literature.

## RESULTS

### MACHINE LEARNING MODELS PERFORMANCE

Among individual classifiers, the SVC model had the highest sensitivity, classifying 90.91% of the pathogenic variants. In contrast, the LR2 algorithm was the one with the highest specificity, classifying 90.21% of the benign variants. The RF algorithm resulted in lower sensitivity and specificity than the two previous models, but was also able to correctly discriminate benign and pathogenic variants. The MLe- KCNQ2 ensemble algorithm improved the overall results of individual models as expected, by correctly classifying 91.11% and 90.21% of pathogenic and benign variants, respectively. AUC-ROC values, above 0.9 in most cases, also suggest that the trained models discriminate excellently between benign and pathogenic variants (Table 1). The values shown in Table 1 represent the average of the results obtained for the train and test sets.

**Table 1.** Performance of trained models. The MLe-KCNQ2 ensemble model performed better, in terms of sensitivity and specificity, than individual Ridge Regression (LR2), Support Vector Classifier (SVC) and Random Forest (RF) classifiers. AUC-ROC values reflect the quality of all models.

| Models | Sensitivity (%) | Specificity (%) | AUC-ROC |
| --- | --- | --- | --- |
| LR2 | 90.67 | 90.21 | 0.91 |
| SVC | 90.91 | 88.00 | 0.90 |
| RF | 87.87 | 87.56 | 0.88 |
| MLe-KCNQ2 | 91.11 | 90.21 | 0.91 |

### RELEVANT FEATURES IN PATHOGENICITY PREDICTION

To determine which variant attributes had the highest influence on pathogenicity prediction; we examined the subset of relevant features previously selected by the feature selection method. This procedure cannot be easily estimated in the ensemble model, as the feature selection is performed in a different way for each individual algorithm of which it is composed. However, an overview of the relevant features in the ensemble model can be obtained by combining the relevant and shared features between the LR2, SVC and RF models

The AlphaFold pLDDT score, the location of the mutation in the FRAG2 of the structural landscape of the K_v_7.2 channel (i.e., residues 195-367) and the MTR score were the top 3 relevant features shared by LR2, SVC and RF models. These attributes together with changes in aromaticity (from non-aromatic to non-aromatic, from aromatic to non-aromatic and from non-aromatic to aromatic), the impact of variants on functional domains such as the voltage-sensing domain or topological domains such as the Cytoplasmic or S4 segments, and the conservation across evolution value across species of the substituted residue constitute the top 10 relevant features for predicting the pathogenicity of variants in K_V_7.2. Taking into account the three subgroups into which the 16 designed features can be classified (See Methods), 2 out of 3 readily available structural information attributes constituted the top 2 relevant features in pathogenicity prediction of missense variants in *KCNQ2*.

### COMPARING MLe-KCNQ2 WITH OTHER PREDICTIVE SOFTWARES

We then compared our novel model with the ability of six *in silico* tools to predict the class of the collected KCNQ2 missense variants. Performances of these tools are shown in Figure 3. FATHMM showed the top score for sensitivity, classifying 99.67% of the collected pathogenic variants. However, the performance of FATHMM was really poor in terms of specificity as it was not able to correctly classify more than 8% of the collected benign variants. Similar results were obtained for the rest of the online software. While pathogenic variants were correctly predicted in CONDEL (99.01%), PolyPhen2 (93.44%), MutPred2 (92.46%), SIFT (83.28%) and PROVEAN (92.79%); none of them was able to identify more than 50% of benign variants. Of note, 50% of the benign variants were incorrectly classified by all these tools. These results clearly indicate that the chosen online predictive software did not correctly discriminate between pathogenic and benign variants in KCNQ2. In contrast, the novel MLe-KCNQ2 algorithm was able to effectively predict the pathogenicity of KCNQ2 variants, by correctly classifying both benign and pathogenic variants with sensitivity and specificity scores above 90.2%.

**Figure 3.**
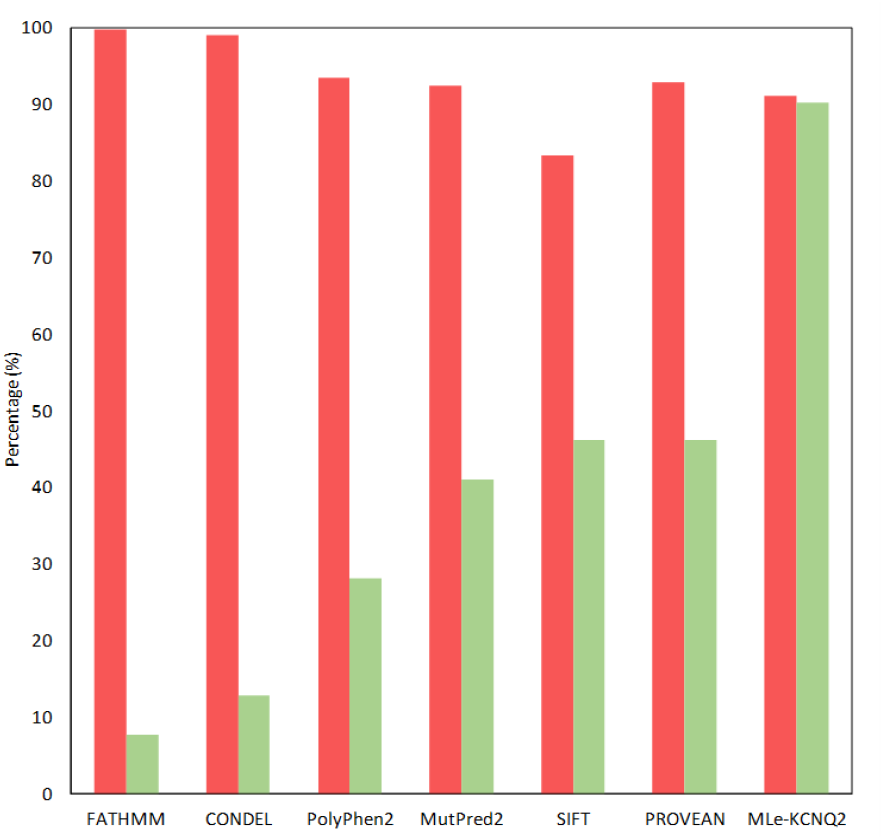
Performance of MLe-KCNQ2 in comparison to other software. Sensitivity (red) and specificity (green) scores are compared using the collected data. The novel MLe-KCNQ2 was the best-performing software by being able to identify both benign and pathogenic variants in KCNQ2 in a similar and efficient manner.

#### The N780T variant has a large prevalence in the human population

In the course of our analysis, we noted that the N780T variant, which is labeled as benign in ClinVar and LOVD3 databases, was classified as such by all these tools. However, the prevalence of this variant is higher than 60% of the population collected in the gnomAD database, which suggest that it represents allelic variant with little or no pathological consequences.

### BIOLOGICAL INTERPRETATION OF SELECTED MLe-KCNQ2 PREDICTIONS

Most confidently predicted benign and pathogenic variants by MLe-KCNQ2 are shown in Figure 4. While variants predicted to be pathogenic (Figure 4B and 4C) were mainly found to affect the VSD domain (R144W, W146R, R201C, I205M, T217I and T217A), in particular the S4 segments and the S2-S3 linker, variants affecting the hTW-hB linker were mostly predicted to be benign (R375C, P379L, G380R, P420L, P420R, P420S, P420Q, A421V and P424S) (Figure 4A and 4C). Variants predicted to be pathogenic were also found in the CaM-interaction domain (D355Y, W360R and Y363C), particularly affecting the intracellular hTW, and in the pore domain (W270C). V22M was a variant predicted with “high confidence” to be benign located at the N-terminus of the K_v_7.2 channel.

**Figure 4.**
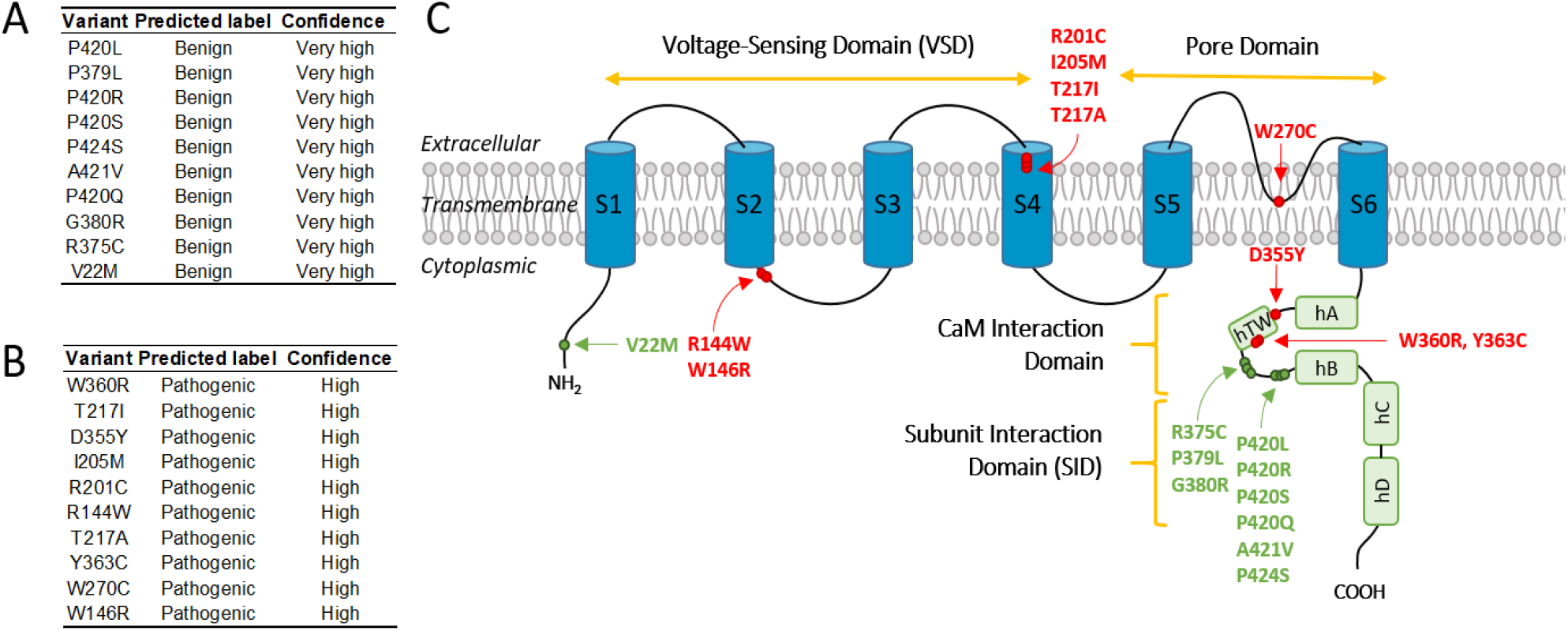
KCNQ2 variants for biological interpretation, their pathogenicity prediction and mapping. **(A)** Most confidently predicted 10 benign and **(B)** top 10 pathogenic predicted variants using MLe-KCNQ2 software. Benign variants have been predicted with a “very high” confidence value (i.e., with a value equal to or greater than 90%); while pathogenic variants have been predicted with a “high” confidence value (i.e., in the range of equal to or greater than 80% and less than 90%). **(C)** Mapping of these variants on the structure of a monomer of the K_v_7.2 channel. The picture depicts the main topological and functional domains of a channel monomer. The six transmembrane segments (S1-S6) are shown in blue and the five intracellular helices in green. The functional voltage-sensor domain (VSD) is composed of segments S1-S4, the pore domain of segments S5-S6, the subunit interaction domain (SID) of helices C-D and the calmodulin interaction domain (CaM) of helices hA, hTW and hB. The location of the benign (green) and pathogenic (red) variants is also depicted.

## DISCUSSION

Accurate pathogenicity prediction of missense variants is of great importance in genetic studies and clinical diagnosis [Qi et al., 2021]. In clinical practice, when a genetic variant is found in the setting of disease, the ACMG (American College of Medical Genetics and Genomics) recommends a variety of techniques to be employed in an attempt to classify the variant as either benign or pathogenic [Richards *et al*., 2015]. However, since the production of genome sequencing data has become routine, the interpretation of the large number of *de novo* discovered variants is posing a major challenge [Lappalainen *et al*., 2019]. *In silico* prediction softwares go some way to address this major issue, but often yield inconsistent results [Chun & Fay, 2009]. ML techniques offer an opportunity to improve pathogenicity prediction tools [Draelos *et al*., 2022].

The structure of a protein is intimately linked to its stability, function and interactions [Ittisoponpisan *et al*., 2019]. It is therefore expected that the structure would be capable of answering a vast number of biological questions including the pathogenicity of variants [Burley *et al*., 2018; Ginalski, 2016]. Despite efforts over many years, only 35% of human proteins map to a PDB entry with the 3D atomic coordinates, and in many cases the structure covers only some fragments of the sequence [Tunyasuvunakool *et al*., 2021; Bateman *et al*., 2021]. For instance, in the structure obtained for Kv7.2 by cryo-EM recently, the spatial location of 523 amino acids (~ 60% of the entire sequence) are not present in the PDB file.

This work pursued two main objectives: (1) developing a ML model to improve accuracy for predicting the pathogenicity of *KCNQ2* missense variants by combining biological-evolutive information, physico-chemical changes and structural features; and (2) creating a framework that can be extended to the prediction of the pathogenicity of variants occurring in other genes. Although there was currently no specific ML based software for *KCNQ2*, similar approaches have been applied to *KCNQ1*, another member of the *KCNQ* family [Phul *et al*., 2022; Draelos *et al*., 2022; Li *et al*., 2017].

First of all, we characterized a set of collected variants combining traditional evolutive features such as evolutive conservation value of the substituted residue; biological information such as mutated residue or affected K_v_7.2 channel domains and physico-chemical changes (i.e., charge, hydrophobicity, molecular weight...); with easily-obtainable genomic and structural information. The incorporation of MTR score and AlphaFold2 pLDDT value constitutes an innovative feature about MLe-KCNQ2 software, which sets it apart from other tools. As a result, a combination of biological-evolutive information, physico-chemical changes and structural features have been considered for pathogenicity prediction. Among the features that influenced the classification of *KCNQ2* variants by MLe-KCNQ2, designed structural descriptors were the most relevant as we expected, followed by the MTR score. The addition of these three novel features in MLe-KCNQ2 allowed increasing the predictive power, resulting in sensitivity and specificity scores of 91.11% and 90.21%, respectively.

We also compared the prediction performances of six *in silico* tools to that of MLe-KCNQ2, and found that the presented gene-specific model outperformed the other softwares. None of the online tools were able to identify benign variants in *KCNQ2* with adequate values (<8-50% in all cases), although pathogenic variants were correctly classified by all of them. Our results suggested that commonly used *in silico* tools could not correctly discriminate between pathogenic and benign variants in *KCNQ2*. We believe that these results can be explained by the type of variants with which each of the algorithms has been trained. While tools like FATHMM, CONDEL, PolyPhen2, MutPred2, SIFT or PROVEAN were trained using information from genome-wide genetic variation data [Pejaver *et al*., 2020; Shihab *et al*., 2013; Choi *et al*., 2012; González-Pérez & López-Bigas, 2011; Adzhubei *et al*., 2010; Ng & Henikoff, 2001]; MLe-KCNQ2 was specifically trained with *KCNQ2* missense variants. This reflects that development of genome-wide tools was based on heterogeneous datasets including a wide range of proteins with diverse functions and associated diseases [Phul *et al*., 2022]. As a direct consequence, their prediction accuracy can vary between genes [Richards *et al*., 2015]. In contrast, MLe-KCNQ2 was trained with specific variants of the *KCNQ2* gene with an emphasis on its particularities, such as the class imbalance scenario between benign and pathogenic variants as well as the lack of experimental structure for the design of 3D-features. From the outset, our software was designed to fit the *KCNQ2* gene problem. Therefore, it makes sense that MLe-KCNQ2 had better classification metrics than these genome-wide tools.

As the results of MLe-KCNQ2 were promising compared to commonly employed tools we decided to give a biological interpretation of a subset of selected variants in order to justify MLe-KCNQ2 predictions and highlight their usefulness. As electrophysiological studies were not available for all the variants, we obtained this information from some experimental studies recently reported in the literature. These works allowed us to justify 3 variants. Firstly, T271I mutation is located in the S4 domain of the K_V_7.2 channel and was predicted as a pathogenic variant. The S4 domain is part of the voltage sensor, has 4 positively charged arginine residues and its movement in response to changes in membrane potential is the basis for the voltage sensitivity of these channels [Sun & MacKinnon, 2017; Cui, 2016; Nakajo & Kubo, 2015]. In addition, experimental studies have shown that mutations in the S4 domain cause significant changes in the activation of K_v_7.2 and would be associated with a pathogenic clinical picture [Zhang *et al*., 2020; Orhan *et al*., 2014]. Recent studies done at the Fudan University Children’s Hospital in China in infants who had seizures in the first month of life determine that this mutation results in encephalopathy [Xu *et al*., 2021]. Secondly, the R144W mutation is located in the S2 transmembrane segment and is therefore part of the voltage-sensing domain. It was also classified as a pathogenic variant by MLe-KCNQ2. Functional studies performed by whole-cell patch-clamp show that activation of homomeric K_v_7.2 R144W/Q/G channels is shifted to the left, suggesting gain-of-function effects (i.e., the probability of opening the gate increases at depolarized potentials) and a pathogenic clinic [Miceli *et al*., 2022]. Finally, the W360R mutation is located in the C-terminal region of the K_v_7.2 channel, specifically in the hTW. MLe-KCNQ2 also predicted W360R as a pathogenic variant with high confidence. Helix TW is located between helices A and B, within the calmodulin (CaM) binding domain of the protein. In addition, the interaction with CaM could be affected in the three-dimensional space due to the substitution of tryptophan (W), non-polar amino acid, by arginine (R), a polar residue. Structurally, this change in polarity could lead to a conformational effect sufficiently relevant for the interaction with CaM to be affected in T359A and, therefore, lead to a pathological clinical scenario. Furthermore, experimental studies have demonstrated the importance of residues extending from T359 to Y362 for CaM binding and it has been suggested that the reduction of this interaction could explain their pathogenicity [Gomis-Perez *et al*., 2015; Richards *et al*., 2004]. In all cases, the predictions made for T217I, R144W and W360R by MLe-KCNQ2 are supported by previous studies.

Finally, in this work we created a framework to develop AI algorithms that can be extended to the prediction of the pathogenicity of variants occurring in other genes. We also demonstrated the extrapolation capability of the working scheme by creating a preliminary model for *KCNH2* gene. The sensitivity and specificity values achieved, higher than 90%, demonstrate that the scheme followed for MLe-KCNQ2 software can be the starting point for the development of other gene-specific predictive software.

## STUDY LIMITATIONS AND FUTURE DIRECTIONS

This work has, among others, four main limitations. The first is that only *KCNQ2* missense variants were included in the design of the MLe-KCNQ2 algorithm. As a direct consequence, MLe-KCNQ2 may not accurately predict neither other *KCNQ2* molecular variants (e.g., frameshift, nonsense...) nor missense mutations in other voltage-gated potassium channels. The second limitation is that MLe-KCNQ2 cannot differentiate between the wide range of phenotypes in the *KCNQ2* encephalopathies. It also cannot determine whether a mutation predicted as pathogenic involves a gain-of-function or a loss-of-function. Pathogenic variants may have a range of effect size and penetrance [Qi et al., 2021]. These difficulties make it necessary to carry out complementary experiments that validate the ML result will help to build trust in approaches and outputs [Vamathevan *et al*, 2019]. This is the main reason why a molecular validation of the predicted variants is highly recommended. The third limitation is the size of the dataset. Although a substantial effort was spent in the compilation of unique *KCNQ2* missense variants, only 353 had a clear or sufficiently reliably clinic description. When larger data sets become available, better predictions could be achieved with MLe-KCNQ2. The last limitation would be the variant reclassifications that regularly occur in clinical databases such as Clinvar [McInnes *et al*., 2021]. If the training data is not updated as these reclassifications occur, any ML-based model would be obsolete and would diminish its clinical relevance.

With available data becoming “larger”, ML algorithms are going to systematically generate improved outputs and new interesting applications are expected to follow [Vamathevan *et al*, 2019]. As a consequence, we suggest further work on the dataset for missense variants of *KCNQ2* by incorporating unseen variants from gnomAD database (https://gnomad.broadinstitute.org/) or recently reported studies. Designing new features for variant characterization such as change in number of hydrogen donor sites or change in number of hydrogen acceptor sites would improve classification metrics as some works remark [Phul *et al*., 2022]. Advances in protein structure prediction (e.g., AlphaFold2) as well as cryo-EM technologies could lead in the design of more complex 3D-features that could make a breakthrough in the prediction of variant pathogenicity. More complex artificial intelligence (AI) methods can also be employed, for example, deep learning techniques. Although there are already a number of studies that have worked with neural networks, one major drawback of deep learning is that it requires an immense amount of data [Jakhar & Kaur, 2019]. Transfer learning offers an opportunity to leverage the power of deep learning in situations where data is limited [McInnes *et al*., 2021]. This emerging approach consists of training a model using data from a well-studied gene (X) and then refine the model with data from a poorly studied gene (Y). The resulting model may perform very well on Y because the “lessons” learned in modeling X transfer well to Y [Taroni *et al*., 2019].

Despite these limitations, MLe-KCNQ2 software represents a major advance in the clinical diagnosis of KCNQ2-related encephalopathies. This work has provided new opportunities in the study of the K_v_7.2 channel that are expected to be further exploited in the future. In addition, the analysis of the pathogenic landscape could help identifying previously unknown domains within the protein sequence, and to discover new functions and possible new pathways for treatments.

## CONCLUSIONS

Here, we provide a powerful tool to predict the pathogenicity of KCNQ2 missense variants. The strength of MLe-KCNQ2 software relies on its performance with both types of variants, benign and pathogenic, with an estimated specificity and sensitivity above 90%. We also found that combining traditional physico-chemical and biological-evolutive features with easily-obtainable structural information could be the key to creating an optimal predictive model. MLe-KCNQ2 will be a helpful tool to improve the accuracy of genetic diagnoses for KCNQ2-related encephalopathies. The framework employed here can be extended to the prediction of the pathogenicity of variants occurring in other genes, as we demonstrated with hERG ion channel.

## SUPPLEMENTARY MATERIAL

Supplementary Material, code, and original data are available at https://github.com/KCNQlab

## Notes

### Competing Interest Statement

The authors have declared no competing interest.

### Summary of Updates

Minor typos in the author names

